# A modular intramolecular triplex photo-switching motif that enables rapid and reversible control of aptamer binding activity

**DOI:** 10.1101/2022.05.22.492975

**Authors:** Tuan Trinh, Ian A. P. Thompson, Finley Clark, Jacob M. Remington, Michael Eisenstein, Jianing Li, H. Tom Soh

**Author notes:** **Corresponding authors: H. Tom Soh** – Department of Radiology and Electrical Engineering, Stanford University, CA, 94305, United States. **email:**., **Jianing Li** – Department of Chemistry, The University of Vermont, Burlington, VT, 05405, United States. **email:**.

## Abstract

DNA switches that can change conformation in response to certain wavelengths of light could enable rapid and non-invasive control of chemical processes for a wide range of applications. However, most current photo-responsive DNA switches are limited either by irreversible switching or reversible switching with impractically slow kinetics. Here, we report the design of an intramolecular triplex photoswitch (TPS) design based on single-stranded DNA that undergoes rapid and reversible photoswitching between folded and unfolded states through isomerization of internal azobenzene modifications. After optimizing the performance of our photoswitch design, we used molecular dynamics (MD) simulations to reveal how individual azobenzenes contribute to the stabilization or destabilization of the triplex depending on their photoisomerization state. By coupling our TPS to an existing aptamer, we can reversibly modulate its binding affinity with less than 15 seconds of UV light exposure. We further demonstrate reproducible shifting in affinity over multiple cycles of UV and blue light irradiation without substantial photobleaching. Given that our TPS can introduce switching functionality to aptamers without manipulating the aptamer sequence itself, we believe our design methodology should offer a versatile means for integrating photo-responsive properties into DNA nanostructures.

## Introduction

Nucleic acids offer a powerful and versatile building block for the construction of nanostructures due to their ease of synthesis and highly specific and programmable molecular recognition.^1^ Recent advances in DNA nanotechnology have enabled the construction of molecular machines that can respond to specific external triggers. These designs allow precise nanoscale control over chemical processes in dynamic materials,^2^ catalytic systems,^3^ and controlled-release drug delivery.^4^ Most mechanisms use chemical ‘fuel’ (*e*.*g*., DNA, redox, pH) to exert stimulus-specific control over DNA nanostructures.^5-10^ However, these approaches require precise and repeated addition of reagents directly into the system, which accumulate over many on-off cycles, leading to contamination of the microenvironment that can potentially limit the lifetime of these devices.^11^ As such, it is highly desirable to design nanostructures that can rapidly and reversibly respond to non-invasive stimuli.

Photoresponsive DNA nanomaterials are an attractive alternative in this regard, as light can be delivered instantaneously and non-invasively, enabling precise temporal and spatial control without the need to add external reagents.^5^ The most common method for designing light-activated nanostructures is through the insertion of photocleavable molecules or caging moieties within the DNA strand.^12-15^ However, these modifications undergo an irreversible structural change upon irradiation, which limits their use in applications when reversibility is required – for example, controlled catch-and-release of biomolecules or on-demand starting and stopping of chemical reactions. Reversible light-based control of DNA nanostructures can be achieved using photo-switchable molecules such as azobenzenes. Azobenzene is a photo-responsive organic molecule that can isomerize from its planar *trans-* form to the non-planar *cis-* form after UV light irradiation (λ = 300–400 nm), and vice versa following exposure to blue light (λ > 400 nm).^16^ For example, DNA duplex hybridization can be controlled through the photo-isomerization of internal azobenzene modifications, where the molecules’ *trans-* and *cis-*conformation stabilizes or destabilizes the duplex, respectively. However, the high stability of DNA duplexes leads to slow isomerization rates and inefficient photoswitching.^17^ For example, Mao *et al*. designed a light-controlled ATP-binding aptamer by incorporating azobenzene moieties within the binding structure, but this design required up to 1 hour of ultraviolet (UV) irradiation to switch the aptamer from a binding to a non-binding state.^18^ These slow switching kinetics are a clear limitation for most applications, where prolonged UV exposure can damage both the DNA nanostructure and the underlying biological system. Liang *et al*. were able to achieve a faster response time with an azobenzene-incorporating DNA intermolecular triplex structure, demonstrating switching within only 10 min of illumination at high temperature (50 °C).^19^ However, their switch design was based on three unlinked DNA strands and thus was only capable of a unidirectional, irreversible response, limiting its utility.

Inspired by the latter work, we report an intramolecular DNA triplex-based motif composed of a single continuous DNA strand that undergoes rapid, reversible azobenzene-mediated photo-switching. Our triplex photo-switch (TPS) construct can be rapidly switched between folded and unfolded states through the isomerization of azobenzene. This allows precise control of the TPS, which opens in response to UV light and closes in response to blue light, with reversibility over many on-off cycles. To understand its switching mechanism, we employed all-atom molecular dynamics (MD) simulations and investigated the structural and energetic landscapes of the TPS construct. Finally, we designed a photo-responsive aptamer switch by coupling the TPS to a well-studied ATP aptamer.^20^ Upon UV exposure, the aptamer’s binding affinity was enhanced by an order of magnitude within approximately 13 seconds, and this effect could be rapidly reversed upon exposure to blue light (λ > 400 nm). This design achieved fast photo-switching response with minimal photobleaching after each cycle. This is a significant improvement over previous work that required multiple minutes of exposure. Finally, unlike the previous designs,^18, 21^ our TPS acts as an independent structural component that does not depend on the aptamer sequence, thus allowing ready integration of photo response properties into wide range of DNA nanostructures. These results demonstrate the potential usefulness of the TPS as a versatile and modular building block for photo-responsive DNA nanotechnology.

## Results and discussion

### Design principle of the triplex photo-switch (TPS)

We hypothesized that by incorporating photoactive azobenzene molecules into one strand of an intramolecular DNA triplex motif, it may be possible to reversibly modulate that triplex structure via UV and blue light exposure. Intramolecular DNA triplexes are formed with three domains; the first and second consist of stretches of homopurines and homopyrimidines, respectively, that form a canonical Watson-Crick duplex, while the third strand consists of homopyrimidines and binds within the major groove of this duplex via non-canonical Hoogsteen base-pairing (**Fig. 1**).^22^ We posited that photo-responsive structure-switching behavior could be introduced into such triplexes by inserting a defined number of azobenzene moieties within the third strand. Briefly, when the azobenzene is in its *trans*-state, the construct should form a stable intramolecular DNA triplex, facilitated by the intercalation of the azobenzene group into the duplex structure. The UV-induced transition to *cis*-azobenzene would then destabilize this structure by eliminating this intercalation and creating steric hindrance that disrupts the Hoogsteen hydrogen bonding of neighboring bases.

**Figure 1.**
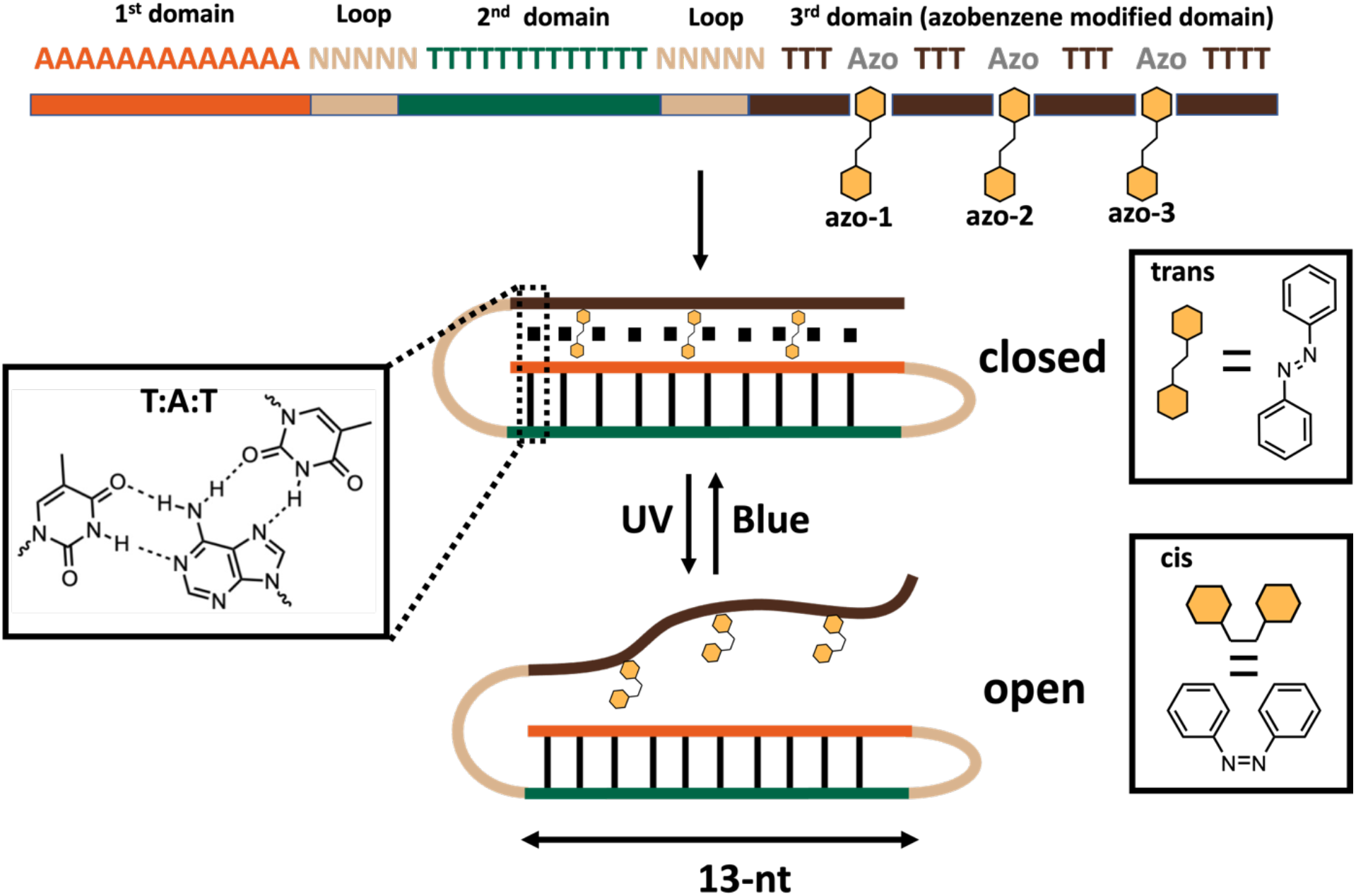
Overall design and working principle of the triplex photo-switch (TPS). The TPS consists of three intertwined helical domains that form an intramolecular DNA triplex. Azobenzene moieties are incorporated into the third domain. UV/blue light exposure induces conformational changes in the azobenzene moieties that cause the TPS to switch between folded and unfolded states. In the *trans* state, azobenzenes participate in stacking and can strengthen triplex formation. After isomerization with UV light, *cis*-azobenzenes can no longer intercalate and sterically distort the third strand, causing it to dissociate. The depicted structure represents the **13-3** TPS.

To test this hypothesis, we designed a triplex comprising two poly-T strands and one poly-A strand. This G-C-free sequence was chosen to ensure triplex folding at physiological pH (∼7.4) and eliminate any pH-dependent effects in folding that occur due to the protonation-dependence of CGC triplex formation.^23^ Each of the three domains consisted of 13 bases, with 5-nt loop sequences connecting the domains together. The two ends of each loop were designed to weakly hybridize to each other through a single base-pairing, thereby promoting intramolecular folding of the structure.

### Tuning triplex stability and folding characterization

We first assessed how the number of azobenzenes introduced into this switch design influences its stability and folding. We synthesized and tested several triplex constructs with zero to four azobenzene moieties inserted into the third, Hoogsteen-paired strand (sequences in **Table 1** and **Supporting Table S1)**. The ionic environment plays an important role in the stability of the overall structure, and we performed our experiments in 1x PBS, pH 7.4 containing 1 mM Mg^2+^ to mimic relevant physiological Na^+^ and Mg^2+^ concentrations (approximately 150 mM and 0.5 to 1 mM, respectively).^24-25^ We used UV-vis spectrophotometry to evaluate the stability of these triplex constructs by assessing melting-induced changes in light absorbance at 260 nm. Folded DNA triplex structures exhibit a biphasic melting curve, wherein the less stable Hoogsteen strand dissociates first, followed by denaturation of the underlying duplex into a random coil (**Supporting Fig. S1**). By monitoring the change in the Hoogsteen melting point (T_m1_), we can gain insights into the triplex stability. Whereas the second, duplex-associated melting point (T_m2_) stayed constant for all four constructs tested, we observed changes in T_m1_ that indicated altered stability of the triplex structure relative to the number of azobenzene groups incorporated (**Table 1, Fig. 2a**). T_m1_ was greatest when a single azobenzene moiety was incorporated into the third strand (T_m1_ = 39–40 °C), suggesting that this single *trans-*azobenzene could intercalate between neighboring bases and thereby stabilize the triplex structure. As the number of azobenzene moieties increased, we observed a gradual decrease in triplex stability, with a ∼1–3°C decrease in T_m1_ per added azobenzene. Interestingly, we observed no triplex formation at room temperature after adding four azobenzenes (**Supporting Fig. S2**). To achieve robust switching between the duplex and triplex conformations of the construct, we aimed to select a triplex design with a T_m1_ only slightly higher than room temperature, which should be fully folded in the *trans-* state but easily destabilized following isomerization to the *cis-* state. We also hypothesized that maximizing the number of azobenzene moieties would increase the probability of isomerizing at least one azobenzene immediately after irradiation, leading to faster switching kinetics.^26^ Based on these criteria, we chose the 13-nt triplex with three azobenzene moieties (termed **13-3**) for further characterization.

**Table 1.**
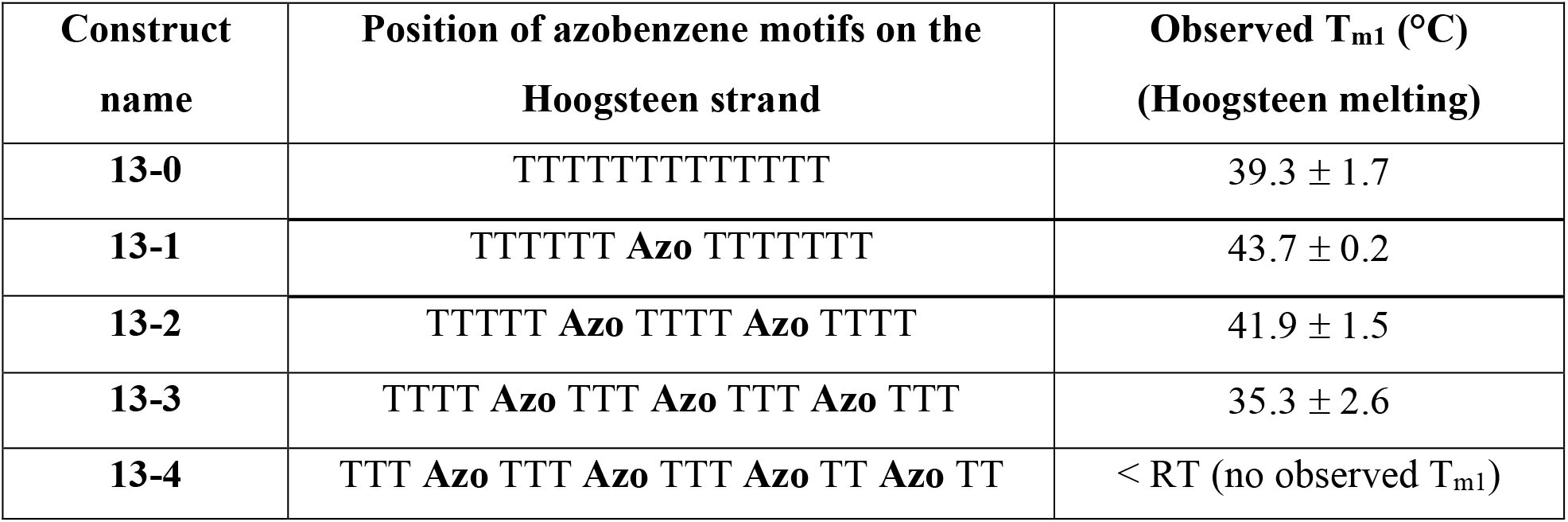
Hoogsteen strand sequences of studied triplex constructs with their observed T_m1_.

**Figure 2.**
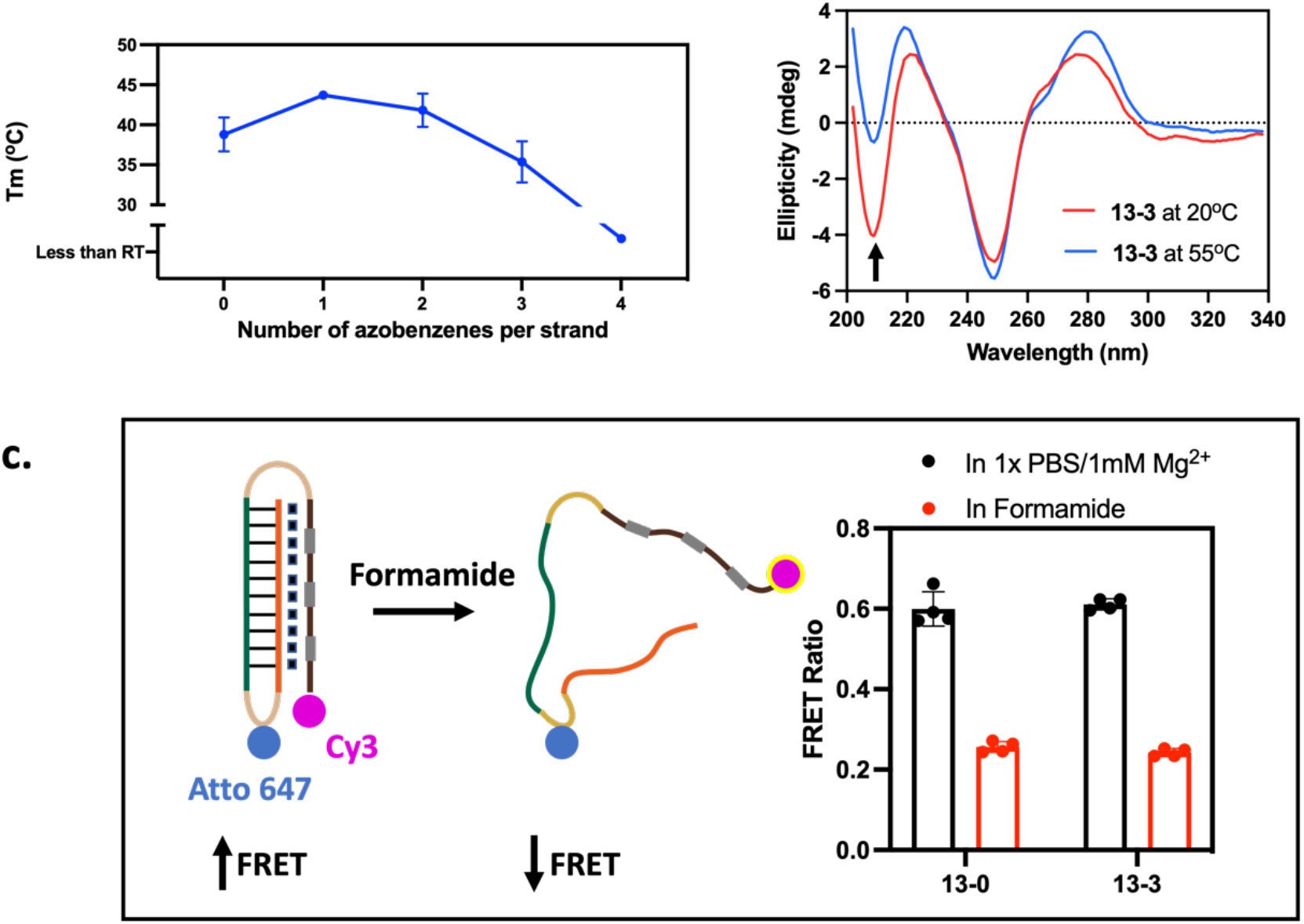
Triplex formation of constructs containing different numbers of azobenzene moieties. **a**) Melting temperature of the Hoogsteen base-paired third strand (T_m1_) in a 13-nt triplex construct with 0–4 azobenzene groups in the third strand. **b)** Circular dichroism (CD) spectra of the **13-3** TPS, which incorporates three azobenzene groups, at 20 °C and 55 °C. The arrow indicates the predicted peak associated with dissociation of the third strand. **c)** A version of **13-3** modified with a pair of dyes to enable monitoring of folding state based on Förster resonance energy transfer (FRET) (left). FRET-based characterization of **13-3** and an azobenzene-free triplex construct (**13-0**) in a physiological buffer (1x PBS/1 mM Mg^2+^) versus after formamide-induced unfolding (right).

To confirm that the folded triplex structure of **13-3** was not distorted by the three azobenzene moieties, we applied circular dichroism (CD) spectroscopy, a sensitive gold-standard technique for studying DNA structural conformation.^27^ Triplex formation produces a characteristic peak at wavelengths of ∼210 nm,^28^ and we confirmed the existence of a triplex structure at 20 °C based on a sharp negative peak at this wavelength (**Fig. 2b)**. This peak was greatly diminished upon heating to 55 °C, a temperature at which Hoogsteen interactions would be disrupted while still preserving Watson-Crick base-pairing (**Supporting Fig. S3**). We then used Förster resonance energy transfer (FRET) ratio to further characterize the folding behavior of **13-3**. We incorporated a Cy3 dye at the 5’ terminus and Atto 643 dye in the first loop domain, such that the two dyes were in close proximity in the fully folded triplex conformation (**Fig. 2c**). This design yielded a strong FRET signal in buffer compared to when the construct was in an unfolded state due to treatment with formamide. These results confirmed that **13-3** forms a properly folded triplex in our buffer conditions. We observed similar folding behavior in a control construct with no azobenzene moieties (**13-0**).

### Characterization of reversible UV-dependent switching

We next tested whether **13-3** undergoes a similar conformational shift in response to stimulation with light. To begin, we used UV-Vis spectrophotometry to monitor the melting profile of the switch before and after UV exposure. After 30 seconds of irradiation with 365-nm UV light, we no longer observed the T_m1_ (Hoogsteen melting) transition that was apparent before light exposure, indicating that this irradiation had effectively disrupted triplex formation (**Fig. 3a**; **Supporting Fig. S4)**. Similarly, when we used CD spectroscopy to compare the conformation of **13-3** before and after UV irradiation, we observed a sharp reduction in the negative peak at 210 nm (**Fig. 3b**), confirming disruption of the triplex structure.

**Figure 3.**
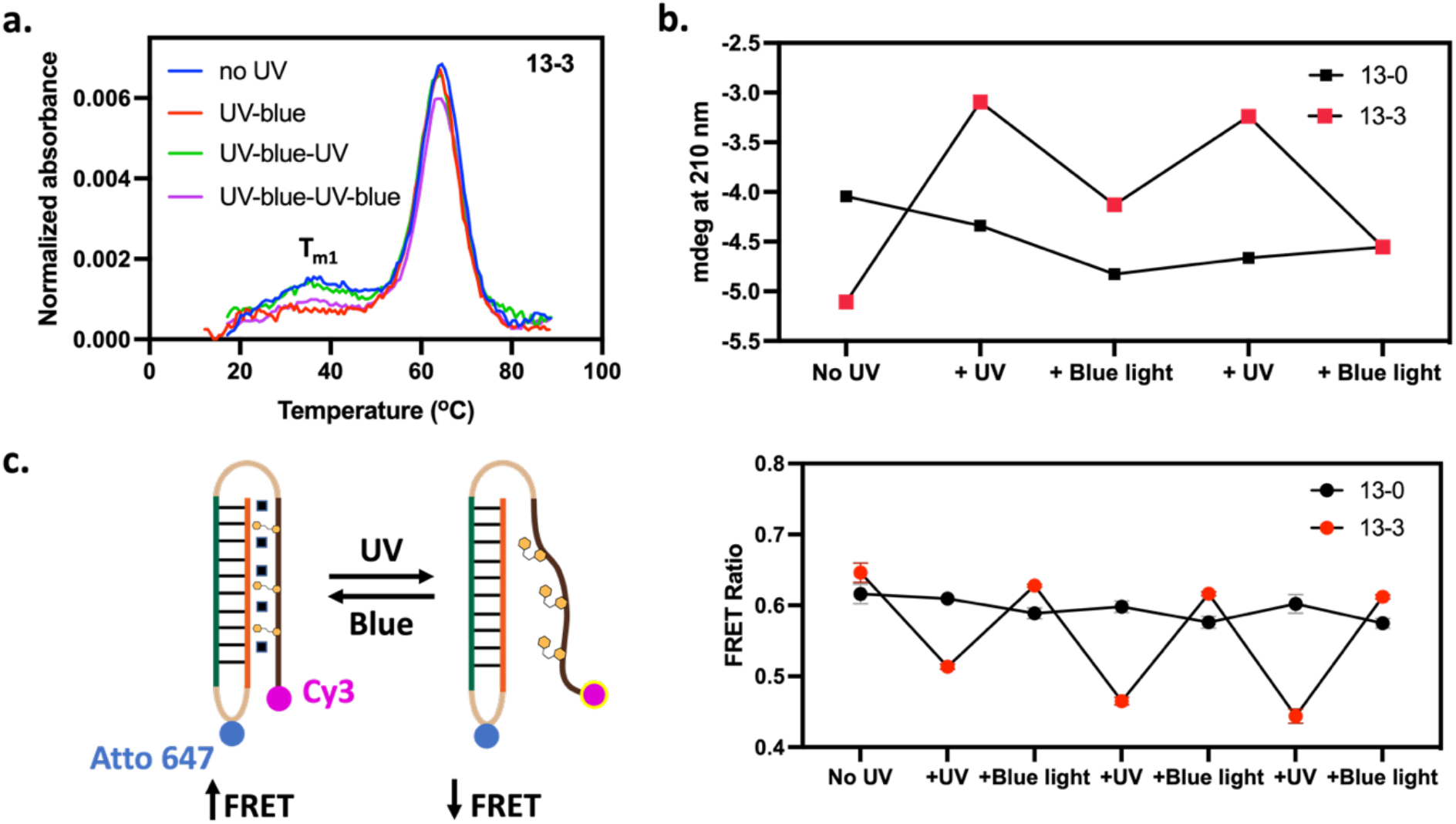
Characterization of TPS switching behavior and reversibility. **a)** First-derivative melting curve of the **13-3** triplex after sequential exposure to UV and blue light. **b)** Amplitude in the ∼210 nm peak from the CD spectra of **13-0** and **13-3** after several rounds of UV and blue light exposure. **c)** FRET characterization of light-induced switching behavior for **13-3** and **13-0** over multiple cycles of UV/blue light illumination.

Importantly, this UV-induced unfolding was fully reversible upon exposing the **13-3** switch to blue light (λ > 400 nm). Both melting curve data (**Fig. 3a; Supporting Fig. S4**) and CD spectra (**Fig. 3b**; **Supporting Fig. S5**) demonstrated that the triplex structure could be fully recovered from its UV-induced unfolded state after just one minute of blue light exposure. In contrast, the azobenzene-free control construct **13-0** showed no switching behavior in response to UV or blue light exposure (**Fig. 3b; Supporting Fig. S6, 7**). We further confirmed the light-induced folding and unfolding of the switch using the FRET-modified version of our **13-3** construct (**Fig. 3c**). We observed a clear drop in the FRET ratio post-UV exposure, indicating an increase in distance between the two dyes relative to the fully folded triple helix, whereas this FRET ratio could be fully recovered after 1 min of blue light exposure. We could repeat this light-induced switching over multiple cycles, whereas control experiments with **13-0** showed no meaningful response to multiple cycles of UV and blue light in the absence of azobenzene moieties. Importantly, we did not observe substantial dye photobleaching after 15 minutes of UV exposure (**Supporting Fig. S8**). These results collectively demonstrate that azobenzene isomerization can selectively trigger unfolding and refolding of the triple helix DNA structure.

### MD simulations provide insight into the structural dynamics of the DNA photo-switches

To gain more insights into the mechanisms underlying these experimental observations, we analyzed the behavior of the three-dimensional (3D) structure of the TPS using all-atom MD simulations. We first simulated the **13-0** and **13-3** constructs for 500 ns at a constant pressure (1 atm) and temperature (27 or 52 °C) in aqueous solution (OPLS3e force field, ∼23,000 atoms with SPC water and counterions for neutralization). We measured the root-mean-square deviation (RMSD) to evaluate the overall flexibility of **13-3** compared to the **13-0** control construct and found that introducing azobenzene reduces the stability of packing and increases the global flexibility of **13-3** compared to **13-0** (**Fig. 4a**). Simulations at higher temperature (52 °C) showed a great increase in the RMSD value of **13-3** due to partial unfolding (**Supporting Fig. S9**), while that of **13-0** remained unchanged throughout the simulation. This decreased stability is consistent with the trend we observed using UV-vis spectrophotometry, wherein the T_m1_ of **13-3** was decreased by approximately 2–4 °C relative to **13-0** (**Fig. 2a, Table 1**). In contrast to RMSD measurements, which provide an understanding of global structural flexibility, per-residue root-mean square fluctuation (RMSF) measurements can be used to analyze how portions of the structure fluctuate, as measured by the average deviation of a residue over time from that residue’s time-averaged position. RMSF analysis showed that the core of the triplex remained stable at 27 °C, whereas the two loops were the most flexible regions, followed by the azobenzene residues **(Fig. 4b)**. Interestingly, the middle azobenzene in **13-3** was the most stable of the three due to its ability to intercalate within neighboring bases (**Supporting Fig. S10**). This is in keeping with the increased T_m1_ observed for **13-1 (Fig. 2a)**, where the subsequent addition of more azobenzenes in **13-2, -3**, or **-4** exhibited a net destabilizing effect by distorting the triplex structure.

**Figure 4.**
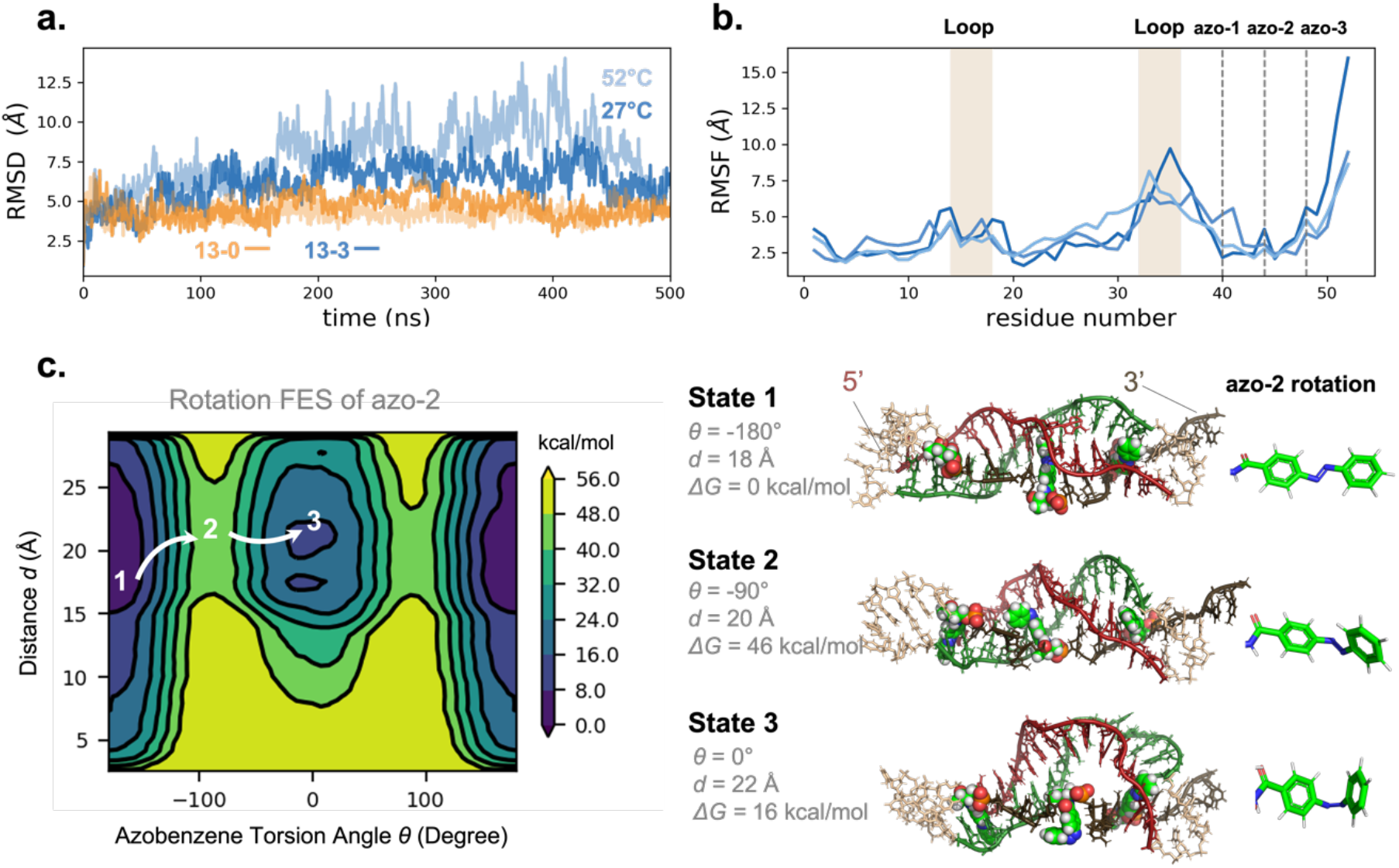
Molecular dynamics (MD) simulations of TPS behavior. **a)** Time evolution of RMSD for **13-0** and **13-3** in 500-ns simulations at 27 and 52 °C. **b)** Per-residue RMSF of three **13-3** simulations at 27 °C show consistent fluctuations. **c)** 2D metadynamics simulations of **13-3** where the middle azobenzene (azo-2) is subject to rotation (as θ), showing changes in the packing of the domains (*i*.*e*., the distance from the azobenzene to the nearest first or second domain nucleotide backbone) and free energy (ΔG). Three representative states (1–3) are labeled, with azobenzenes depicted as spheres in the cartoon and the corresponding conformations as sticks on the right.

Finally, we carried out 2D metadynamics simulations for each azobenzene in **13-3** to study the thermodynamic impact of the rotation of the azo bond and the packing of the third domain relative to the duplex, which was measured as the backbone distance from the azobenzene to the nearest residue in either the first or second domain.^29^ In our simulations, we used a Gaussian biasing force to enhance sampling of the azo bond rotation of each individual azobenzene, with the other two azobenzenes fixed in the *trans*-conformation. Upon convergence, we studied their free energy (ΔG) over the full range of torsion angles spanning from 180° (*trans-*) to 0° (*cis-*). The second azobenzene (**azo-2**) had the highest energetic barrier (45.2 ± 2.1 kcal/mol) compared with **azo-1** (44.0 ± 1.4 kcal/mol) and **azo-3** (42.9 ± 1.5 kcal/mol) (**Supporting Fig. S11**). This is consistent with the heightened stability of that central azobenzene **(azo-2)** revealed by RMSD and RMSF measurements and UV-vis experimental data. Importantly, when rotating only **azo-2**, ΔG for **13-3** is about 16 kcal/mol lower (or more stable) when that azobenzene is in the *trans*-versus the *cis*-conformation (**Fig. 4c**, states 1 and 3). This simulated ΔG estimate is substantially higher than previously-reported ΔG measurements of *trans*-to-*cis* isomerization of free azobenzene in the absence of a triplex structure (10-12 kcal/mol).^30^ This difference can be attributed to changes in ΔG of the full triplex structure, which experiences destabilization through the disruption of azobenzene stacking and triplex structure steric effects and agrees well with our experimental findings in which azobenzene photoisomerization disrupts folding of the third strand.

### Designing a photo-responsive aptamer switch using the TPS motif

We hypothesized that our TPS could be readily integrated into more complex DNA nanodevices to achieve light-activated molecular control, and tested this approach by coupling the TPS with an ATP aptamer that has been widely studied in the literature.^20^ Briefly, we inserted the **13-3** TPS into a linker domain connecting the ATP aptamer and a displacement strand that competitively binds the aptamer sequence (**Fig. 5a**). Based on previous work on tuning aptamer switches,^31^ we determined that an 8-nt displacement strand would achieve robust intramolecular switching while still allowing the coupled aptamer to retain moderate (in this case, ∼1 mM) ligand-binding affinity. We inserted short spacers of 4–5-nt between the aptamer, triplex switch, and displacement strand to minimize the formation of unintended secondary structure between the three domains. Finally, the resulting **Apt-13-3** construct was functionalized with Atto 643 and Cy3 dyes at the 5’ and 3’ ends, respectively, to enable FRET-based measurement of ATP binding and aptamer switching.

**Figure 5.**
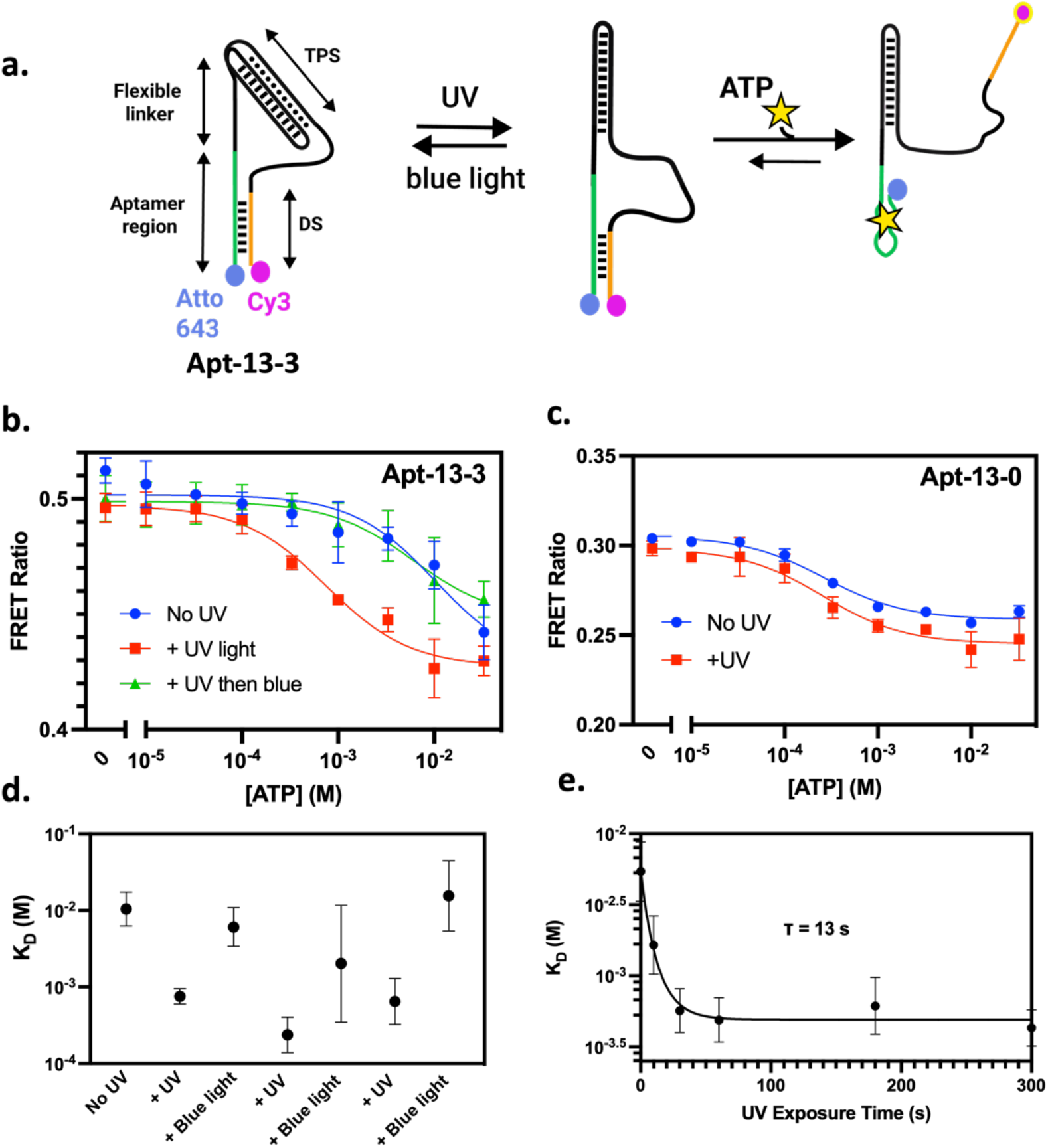
Employing the 13-3 TPS for light-induced modulation of ATP aptamer binding affinity. **a)** Schematic of the **Apt-13-3** construct and its predicted photo-switching behavior, wherein UV irradiation is expected to greatly improve ATP binding by triggering triplex unfolding and subsequent destabilization of the aptamer-bound displacement strand. **b)** FRET-based binding curves of **Apt-13-3** show a reversible 10-fold increase in binding affinity upon UV exposure. **c)** No such switching is apparent with control construct **Apt-13-0. d)** Measuring the shift in ATP binding affinity for **Apt-13-3** over three cycles of UV/blue light-induced switching. **e)** Kinetics of the binding affinity shift for **Apt-13-3** upon UV exposure.

In this design, light-induced conformational changes in the triplex linker are predicted to modulate competitive binding of the displacement strand and thereby influence aptamer folding and target binding.^32^ Prior to UV exposure, the TPS domain adopts a folded triplex state and acts as a compact linker between the aptamer and displacement strand, producing confinement that leads to strong intramolecular competition, resulting in low aptamer binding affinity (high *K*_*D*_). Upon UV light excitation, the triplex structure is disrupted, and the motif forms an extended and more flexible linker, leading to weaker displacement strand competition and increased aptamer binding affinity (low *K*_*D*_).

To evaluate the photo-responsive properties of this switch construct, we measured its ATP binding affinity through a FRET-based binding assay, comparing its binding properties before or after excitation with UV or blue light. As the concentration of ATP increases, competitive ATP binding causes displacement of the Cy3-labeled displacement strand domain within the switch, which leads to decreased FRET signal. We found that aptamer affinity was robustly and reversibly increased ∼10-fold upon UV exposure (**Fig. 5b**). Before illumination, the baseline affinity was K_D_ ∼10 mM (95% CI [2.5-56.3 mM]), which was reduced to ∼700 μM after 30 s of UV excitation (95% CI [0.5-1.3 mM]); this affinity could then be reset back to ∼10 mM after exposure to blue light for 1 minute. In contrast, the same switch construct without azobenzene molecules (**Apt-13-0**) showed no UV-induced shift in affinity (**Fig. 5c**). Importantly, this shift in K_D_ could be induced consistently over several cycles of UV/blue light exposure **(Fig. 5d**). Finally, we quantified the kinetics of UV-induced switching by measuring the binding affinity over a wide range of UV illumination times and found the time constant of photo-responsive activation (*τ*) to be 13 seconds (95% CI [7-24 s] (**Fig. 5e)**. Interestingly, our approach uses TPS as an independent structure motif that does not depend on the aptamer sequence, thus could be adapted to incorporate photo-responsive properties into a wide range of DNA nanostructures.

## Conclusion

In this work, we have designed and characterized an azobenzene-modified photo-responsive switch based on a single-stranded intramolecular DNA triplex structure that can rapidly and reversibly respond to different wavelengths of light. By integrating azobenzene modifications within the triplex Hoogsteen strand, we were able to photo-regulate folding between the opened and closed state of our TPS motif with short-duration UV or blue light exposure. Importantly, we showed that the TPS can undergo this structural change within 13 seconds and reversibly over many on/off cycles. This is in striking contrast to previous designs for photo-responsive DNA materials based on DNA duplexes, which have suffered from inefficient switching and slow kinetics. This is attributable to the low quantum yield of azobenzene in a double-stranded context, which is six times lower than that of azobenzene in a single-stranded context and nine times lower than that of free azobenzene.^17^ By tuning the stability of our intramolecular triplex construct so that T_m1_ is only marginally higher than room temperature, we were able to achieve more rapid photoswitching. Finally, we integrated the TPS motif into an ATP-binding aptamer and demonstrated robust photoregulated activation and inhibition of binding function, with a rapid and reversible 10-fold shift in binding affinity upon UV or blue light exposure. Among other assets, this design offers the advantages of enabling robust, repeated cycling while only requiring short excitation times, minimizing the risk of UV-induced photodamage or bleaching. Since this strategy uses the TPS as an added component to the aptamer structure rather than directly modifying the aptamer itself, the same approach may be amenable to engineering a wider range of photo-responsive aptamers. As such, we anticipate that this or similar constructs should be applicable for achieving light-mediated control of molecular nanodevices in the context of self-assembly, cascade reactions, controlled-release drug delivery, real-time diagnostics, and a range of other applications.

## Methods

### UV-Vis measurement

UV-Vis experiments were performed with a Cary 300 Bio instrument from Agilent Technology with temperature controller. Briefly, a 500 μL sample of DNA in 1 μM in 1x PBS / 1 mM Mg^2+^ (pH 7.4) buffer was annealed from 95 to 4 °C over the course of 1 hour before collecting UV-Vis measurements. The absorbance at 260 nm was monitored as the temperature was increased from 25 to 95 °C at increments of 1 °C/min. The melting temperature (T_m_) was determined from the highest values of the first derivatives.

### CD measurement

CD spectrometry was performed using a 1-mm path-length quartz cuvette on a Jasco-815 spectropolarimeter equipped with a xenon lamp, a Peltier temperature control unit, and a water recirculator. A 300 μL sample containing 5 μM TPS DNA was prepared in 1x PBS/1 mM Mg^2+^ (pH 7.4) buffer. The sample was annealed from 95 °C to 4 °C over the course of 1 hour before collecting CD measurements. CD spectra were obtained (200–340 nm range, 100 nm/min scan rate, 1 nm bandwidth, 3 accumulations) at 20 °C.

### FRET assay

Both **13-0** and **13-3** strands were first functionalized with a Cy3 dye at the 3’ terminus (using solid-phase synthesis) and Atto 643 dye in the first loop domain (through amino-N-hydroxysuccinimide (NHS) conjugation) before the assay, as shown in **Fig. 2c**.

- For the experiment shown in **Fig. 2c**: Two separate **13-3** samples (200 μL) were prepared at 100 nM final concentration. The first sample was prepared in 1x PBS, pH 7.4 with 1 mM Mg^2+^ buffer, then annealed from 95 °C to 4 °C over the course of 1 hour. The second sample was prepared in a solution of formamide: water (90:10 v/v). Both samples were exposed to 3 mins of UV exposure using quartz cuvettes (λ = 365 nm, UV intensity ∼144-187 mW/cm^2^). Following the UV exposure, both samples were allowed to equilibrate for 5 minutes to room temperature. Fluorescence was measured on a Synergy H1 microplate reader (BioTeK) with Cy3 excitation (filter properties: 540 nm center wavelength, 25 nm bandpass width) and measurement of Cy3 emission (590 nm center wavelength, 35 nm bandpass width) as well as Cy3-Cy5 FRET emission (680 nm center wavelength, 30 nm bandpass width). All measurements were taken in triplicate. The FRET ratio for each replicate was calculated as:

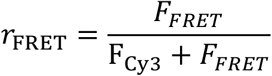
- For the experiment shown in **Fig. 3c**: **13-3** sample was prepared at 100 nM in 1x PBS, pH 7.4 with 1 mM Mg^2+^ buffer, and then annealed from 95 °C to 4 °C over the course of 1 hour. The sample was exposed to alternating cycles of 30 s UV exposure and 1 min blue light in a quartz cuvette. The fluorescence intensity was measured, and the FRET ratio was then calculated using the equation as described above.

### Binding affinity measurements

All aptamer binding measurements were carried out in 1x PBS, pH 7.4, with 1 mM Mg^2+^. Aptamer stocks were first diluted to 100 nM in buffer, then annealed from 95 °C to 4 °C over the course of 1 hour. The aptamer stocks were exposed to 0–3 alternating cycles of 30 s UV exposure and 1 min blue light exposure in a quartz cuvette for the reversibility experiments, or to varying durations of UV light exposure for the aptamer kinetics experiment (UV light intensity = ∼144-187 mW/cm^2^). Following UV exposure during each cycle, the samples were allowed to equilibrate for 5 minutes at room temperature. 20 μL of 100 nM aptamer stock was mixed with 20 μL of ATP at various concentrations in assay buffer to reach final concentrations spanning 0–33 mM ATP. These ATP-aptamer mixtures were incubated at room temperature for 30 min. Fluorescence was measured on a Synergy H1 microplate reader (BioTeK) with Cy3 excitation (filter properties: 540 nm center wavelength, 25 nm bandpass width) and measurement of Cy3 emission (590 nm center wavelength, 35 nm bandpass width) as well as Cy3-Cy5 FRET emission (680 nm center wavelength, 30 nm bandpass width). All measurements were taken in triplicate. Additionally, measurements were performed for a 40 μL sample of assay buffer to determine background signal for later subtraction.

### Analysis of UV/blue light excitation-dependent aptamer binding affinity

All data analysis was performed in Microsoft Excel and GraphPad Prism 8.0.2. First, raw triplicate Cy3 emission (*F*_Cy3_) and Cy3-Cy5 FRET emission (*F*_*FRET*_) vs. ATP concentration ([ATP]) data were processed by subtracting the background values obtained from buffer alone. Then, the FRET ratio for each replicate was calculated as:

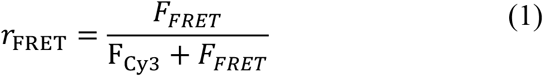

To determine the binding affinity from each combination of aptamer sequence and UV/blue exposure conditions, the FRET data were fitted to a Langmuir isotherm of the form:

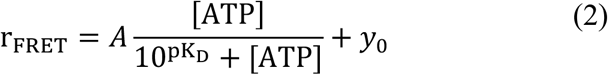

Where pK_D_ = log_10_(*K*_*D*_) was used to extract the binding affinity (*K*_*D*_) of the construct after exposure, *y*_0_ is the background FRET ratio from the construct in the absence of ATP, and *A* is the range of FRET ratio change upon ATP-induced switching. Binding affinities are reported as the best fit value with 95% confidence interval upper and lower bounds from fitting.

To determine the kinetics of the aptamer photo-response, we analyzed the extracted *K*_*D*_ vs UV exposure time data from our binding curve fitting. We fitted this affinity vs. exposure time data using a single-phase exponential decay model of the form:

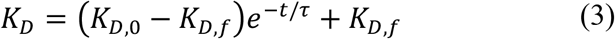

Where *K*_*D*,0_ and *K*_*D,f*_ are respectively the aptamer binding affinities before and after prolonged UV exposure, and *τ* is the time constant that characterizes the kinetics of aptamer photo-switching.

### Molecular modeling

Our models were constructed visualized in Maestro (Schrödinger, Inc.) with help of our in-house program DNA BACon (Building, Assembly, and Construct). All the simulations were performed with the OPLS3e force field with the SPC water model in the Desmond-GPU program (Schrödinger, Inc.), and analyzed with in-house Python and Tcl scripts.

## Author information

### Authors

- **Tuan Trinh** – *Department of Radiology, Stanford University, CA, 94305, United States*.
- **Ian A.P. Thompson** – *Department of Electrical Engineering, Stanford University, CA, 94305, United States*.
- **Finley Clark** – *Department of Chemistry, The University of Vermont, Burlington, VT, 05405, United States*.
- **Jacob M. Remington** – *Department of Chemistry, The University of Vermont, Burlington, VT, 05405, United States*.
- **Michael Eisenstein** – *Department of Radiology, Stanford University, CA, 94305, United States*.

## Data availability

All study data are included in the article and/or supporting information.

## Acknowledgements

This work was supported by the Chan-Zuckerberg Biohub, the Helmsley Trust, W.L Gore and Associates, and the National Institutes of Health (NIH, OT2OD025342). I.A.P.T. would like to acknowledge the support of the Medtronic Foundation Stanford Graduate Fellowship and the Natural Sciences and Engineering Research Council of Canada (NSERC, 416353855). F.C. and J.M.R. were partially supported by an NIH R01 award (R01GM129431 to J.L.) and J.L. was partially supported by the NSF CAREER award (CHE-1945394). The authors would like to thank Prof. Severin T. Schneebeli (University of Vermont) for the helpful discussions, Prof. Eric T. Kool and Prof. Possu Huang (Stanford University) for the use of UV-Vis and CD instruments, respectively.

## Notes

The authors declare no competing financial interest.

## References

1. Seeman, N. C.,, Sleiman, H. F., DNA nanotechnology. Nature Reviews Materials 2017, 3 (1), 17068.

2. Ryssy, J.; Natarajan, A. K.; Wang, J.; Lehtonen, A. J.; Nguyen, M.-K.; Klajn, R.; Kuzyk, A., Light-Responsive Dynamic DNA-Origami-Based Plasmonic Assemblies. Angewandte Chemie International Edition 2021, 60 (11), 5859–5863.

3. Wang, S.; Yue, L.; Li, Z.-Y.; Zhang, J.; Tian, H.; Willner, I., Light-Induced Reversible Reconfiguration of DNA-Based Constitutional Dynamic Networks: Application to Switchable Catalysis. Angewandte Chemie International Edition 2018, 57 (27), 8105–8109.

4. Karimi, M.; Sahandi Zangabad, P.; Baghaee-Ravari, S.; Ghazadeh, M.; Mirshekari, H.; Hamblin, M. R., Smart Nanostructures for Cargo Delivery: Uncaging and Activating by Light. Journal of the American Chemical Society 2017, 139 (13), 4584–4610.

5. Liu, X.; Lu, C.-H.; Willner, I., Switchable Reconfiguration of Nucleic Acid Nanostructures by Stimuli-Responsive DNA Machines. Accounts of Chemical Research 2014, 47 (6), 1673–1680.

6. Idili, A.; Vallée-Bélisle, A.; Ricci, F., Programmable pH-Triggered DNA Nanoswitches. Journal of the American Chemical Society 2014, 136 (16), 5836–5839.

7. Farag, N.; Mattossovich, R.; Merlo, R.; Nierzwicki, Ł.; Palermo, G.; Porchetta, A.; Perugino, G.; Ricci, F., Folding-upon-Repair DNA Nanoswitches for Monitoring the Activity of DNA Repair Enzymes. Angewandte Chemie International Edition 2021, 60 (13), 7283–7289.

8. Prinzen, A. L.; Saliba, D.; Hennecker, C.; Trinh, T.; Mittermaier, A.; Sleiman, H. F., Amplified Self-Immolative Release of Small Molecules by Spatial Isolation of Reactive Groups on DNA-Minimal Architectures. Angewandte Chemie International Edition 2020, 59 (31), 12900–12908.

9. Yurke, B.; Turberfield, A. J.; Mills, A. P.; Simmel, F. C.; Neumann, J. L., A DNA-fuelled molecular machine made of DNA. Nature 2000, 406 (6796), 605–608.

10. Liu, M.; Fu, J.; Hejesen, C.; Yang, Y.; Woodbury, N. W.; Gothelf, K.; Liu, Y.; Yan, H., A DNA tweezer-actuated enzyme nanoreactor. Nature Communications 2013, 4 (1), 2127.

11. Simmel, F. C.; Dittmer, W. U., DNA Nanodevices. Small 2005, 1 (3), 284–299.

12. Young, D. D.; Lively, M. O.; Deiters, A., Activation and Deactivation of DNAzyme and Antisense Function with Light for the Photochemical Regulation of Gene Expression in Mammalian Cells. Journal of the American Chemical Society 2010, 132 (17), 6183–6193.

13. Tam, D. Y.; Dai, Z.; Chan, M. S.; Liu, L. S.; Cheung, M. C.; Bolze, F.; Tin, C.; Lo, P. K., A Reversible DNA Logic Gate Platform Operated by One- and Two-Photon Excitations. Angewandte Chemie International Edition 2016, 55 (1), 164–168.

14. Chandrasekaran, A. R.; Abraham Punnoose, J.; Valsangkar, V.; Sheng, J.; Halvorsen, K., Integration of a photocleavable element into DNA nanoswitches. Chemical Communications 2019, 55 (46), 6587–6590.

15. Liu, M.; Jiang, S.; Loza, O.; Fahmi, N. E.; Šulc, P.; Stephanopoulos, N., Rapid Photoactuation of a DNA Nanostructure using an Internal Photocaged Trigger Strand. Angewandte Chemie International Edition 2018, 57 (30), 9341–9345.

16. Griffiths, J., II. Photochemistry of azobenzene and its derivatives. Chemical Society Reviews 1972, 1 (4), 481–493.

17. Yan, Y.; Wang, X.; Chen, J. I. L.; Ginger, D. S., Photoisomerization Quantum Yield of Azobenzene-Modified DNA Depends on Local Sequence. Journal of the American Chemical Society 2013, 135 (22), 8382–8387.

18. Zhang, X.; Song, C.; Yang, K.; Hong, W.; Lu, Y.; Yu, P.; Mao, L., Photoinduced Regeneration of an Aptamer-Based Electrochemical Sensor for Sensitively Detecting Adenosine Triphosphate. Analytical Chemistry 2018, 90 (8), 4968–4971.

19. Liang, X.; Asanuma, H.; Komiyama, M., Photoregulation of DNA Triplex Formation by Azobenzene. Journal of the American Chemical Society 2002, 124 (9), 1877–1883.

20. Huizenga, D. E.; Szostak, J. W., A DNA Aptamer That Binds Adenosine and ATP. Biochemistry 1995, 34 (2), 656–665.

21. Zhang, L.; Zhang, X.; Feng, P.; Han, Q.; Liu, W.; Lu, Y.; Song, C.; Li, F., Photodriven Regeneration of G-Quadruplex Aptasensor for Sensitively Detecting Thrombin. Analytical Chemistry 2020, 92 (11), 7419–7424.

22. Nikolova, E. N.; Zhou, H.; Gottardo, F. L.; Alvey, H. S.; Kimsey, I. J.; Al-Hashimi, H. M., A historical account of hoogsteen base-pairs in duplex DNA. Biopolymers 2013, 99 (12), 955–968.

23. Amodio, A.; Zhao, B.; Porchetta, A.; Idili, A.; Castronovo, M.; Fan, C.; Ricci, F., Rational Design of pH-Controlled DNA Strand Displacement. Journal of the American Chemical Society 2014, 136 (47), 16469–16472.

24. Melkikh, A. V.; Sutormina, M. I., Model of active transport of ions in cardiac cell. Journal of Theoretical Biology 2008, 252 (2), 247–254.

25. Heikenfeld, J.; Jajack, A.; Feldman, B.; Granger, S. W.; Gaitonde, S.; Begtrup, G.; Katchman, B. A., Accessing analytes in biofluids for peripheral biochemical monitoring. Nature Biotechnology 2019, 37 (4), 407–419.

26. Asanuma, H.; Liang, X.; Nishioka, H.; Matsunaga, D.; Liu, M.; Komiyama, M., Synthesis of azobenzene-tethered DNA for reversible photo-regulation of DNA functions: hybridization and transcription. Nature Protocols 2007, 2 (1), 203–212.

27. Gray, D. M.; Ratliff, R. L.; Vaughan, M. R., [19] Circular dichroism spectroscopy of DNA. In Methods in Enzymology, Academic Press: 1992; Vol. 211, pp 389–406.

28. Gray, D. M.; Hung, S.-H.; Johnson, K. H., [3] Absorption and circular dichroism spectroscopy of nucleic acid duplexes and triplexes. In Methods in Enzymology, Academic Press: 1995; Vol. 246, pp 19–34.

29. Liao, C.; Esai Selvan, M.; Zhao, J.; Slimovitch, J. L.; Schneebeli, S. T.; Shelley, M.; Shelley, J. C.; Li, J., Melittin Aggregation in Aqueous Solutions: Insight from Molecular Dynamics Simulations. The Journal of Physical Chemistry B 2015, 119 (33), 10390–10398.

30. Beharry, A. A.; Woolley, G. A., Azobenzene photoswitches for biomolecules. Chemical Society Reviews 2011, 40 (8), 4422–4437.

31. Wilson, B. D.; Hariri, A. A.; Thompson, I. A. P.; Eisenstein, M.; Soh, H. T., Independent control of the thermodynamic and kinetic properties of aptamer switches. Nature Communications 2019, 10 (1), 5079.

32. Thompson, I. A. P.; Zheng, L.; Eisenstein, M.; Soh, H. T., Rational design of aptamer switches with programmable pH response. Nature Communications 2020, 11 (1), 2946.

