## Supporting Information for "A modular intramolecular triplex photo-switching motif that enables rapid and reversible control of aptamer binding activity"

*Corresponding authors:

**Supporting Information**

**General:** Unless otherwise stated, all commercial reagents and solvents were used without additional purification. 1M magnesium chloride solution, GelStar nucleic acid stain and 100 mM ATP solution were purchased from Thermo Fisher Scientific. Sequagel concentrate, buffer and diluent for denaturing polyacrylamide gel electrophoresis (PAGE) were purchased from National Diagnostics. Atto 643-NHS ester was purchased from ATTO-TEC GmbH. Formamide was purchased from Acros Organic. Sephadex G25 columns, Azobenzene phosphoramidite, cyanine 3 phosphoramidite, and 5'-amino-modifier C6 were purchased from Glen Research. 1x TBE buffer is composed of 90 mM Tris, 90 mM boric acid and 2 mM EDTA with a pH ~8.3 All DNA sequences are listed in **Supporting Table S1**.

**Instrumentation:** Standard automated oligonucleotide solid-phase synthesis was performed on an Expedite 8909 synthesizer from Biolytic Lab Performance. UV-Vis experiments were performed on a Cary 300 instrument from Agilent. DNA quantification measurements were performed by UV absorbance at 260 nm with the NanoDrop 2000 spectrophotometer from Thermo Fisher Scientific. Gel electrophoresis experiments were carried out on a 20 x 20 cm vertical Hoefer 600 electrophoresis unit. Gel images were captured using a ChemiDoc MP System from Bio-Rad Laboratories. Thermal annealing of all DNA structures was conducted using an Eppendorf Mastercycler 96 well thermocycler. Fluorescent binding assays were performed on a Synergy H1 microplate reader (BioTeK).

**Synthesis and purification of azobenzene-modified DNA strands:** DNAs were synthesized using an Expedite 8909 DNA synthesizer at 1 μmol scale. Coupling efficiency was monitored after removal of the DMT 5-OH protecting group. Azobenzene phosphoramidite's coupling time was extended to 15 mins per coupling for better yield. After deprotection with a 1:1 aqueous ammonium hydroxide:methyl amine (AMA) solution for 2 hours at room temperature, the crude product was isolated, dried, and re-suspended in 1:1 (v/v) 8 M urea before loading onto a 15% denaturing polyacrylamide gel (PAGE). The gel was run at 300 V for 4 hours in 1x TBE and then imaged and excised on a TLC plate under a UV lamp. The cut band was crushed and soaked in miliQ water overnight. The solution was then dried to approximately 1 mL before loading onto a

Sephadex G-25 column for desalting. The purified DNA was quantified based on absorbance at 260 nm.

**Model construction.** The **13-3** photoswitch was created based on the template of a small DNA triplex (PDBID: 1BWG). This structure was chosen because one of the canonically base-pairing strands is composed entirely of purines, while the other is composed entirely of pyrimidines. The third strand, composed entirely of pyrimidines, pairs with the purine strand through Hoogsteen interactions. Assuming that our folded photoswitch would retain this structure, we first used the PyMOL program to mutate bases in the template from cytosine to thymine and guanine to adenine, and then converted the appropriate nucleotides in the third strand to a *trans*-azobenzene moiety using the build tools in Maestro (Schrödinger Inc.). The initial models were refined in multiple stages. We first adjusted the nucleotides to maximize contacts via Hoogsteen base-pairing and parallel pi stacking. Then, we performed energy minimization and/or MD equilibration followed by short, unbiased MD simulations (~30 ns and shorter) in Desmond (GPU version, Schrödinger Inc.). When a stable triplex model was achieved, it was subjected to a 100 ns MD simulation with restraints on nitrogen-carbon torsion in the azobenzene to improve the rigidity of the planar conjugated azobenzene. Constructs **13-0, 13-1**, and **13-2** were created in a similar fashion.

**Simulation setup and analysis.** All unbiased and metadynamics simulations used the OPLS3e force field with the SPC water model and were run with Desmond software. Each construct (~23,000 atoms) went through a multi-stage minimization and equilibration procedure, and production runs in the NPT ensemble (27 or 52 °C, 1 bar, Nosé-Hoover thermostat with 1 ps relaxation time, and Martyna-Tobias-Klein isotropic barostat with 2 ps relaxation time) using 2 fs timesteps. The particle mesh Ewald (PME) technique was used for the electrostatic calculations. A van der Waals and short-range electrostatic cutoff of 9 Å was selected and updated with a time step of 2 fs, while long-range electrostatics were calculated every 6 fs. The unbiased simulations were performed for 200 or 500 ns, while the metadynamics simulations were performed for 500 ns. Visualization was performed using VMD, Maestro, and PyMOL. All analysis was performed using in-house Tcl and Python scripts.

**Supporting Table S1 |** DNA sequences used in this work. **Azo** = azobenzene phosphoramidite, **$NH_2$ =** Amino C6 modifier.

| Name | Sequence (5’ → 3’) |
|---|---|
| **13-0** | TTTTTTTTTTTTTGTTTCTTTTTTTTTTTTTATTTAAAAAAAAAAAAA |
| **13-1** | TTTTTT/**Azo**/TTTTTTTGTTTCTTTTTTTTTTTTTATTTAAAAAAAAAAAAA |
| **13-2** | TTTTT/**Azo/**TTTT/**Azo**/TTTTGTTTCTTTTTTTTTTTTTATTTAAAAAAAAAAAAA |
| **13-3** | TTTT/**Azo**/TTT/**Azo/**TT/**Azo**/TTTGTTTCTTTTTTTTTTTTTATTTTAAAAAAAAAAAAA |
| **13-3 functionalized with Cy3 and Atto-643** | **/Cy3**/TTTT/**Azo**/TTT/**Azo/**TT/**Azo**/TTTGTTTCTTTTTTTTTTTTTAT/**Atto643/**TTAAAAAAAAAAAAA |
| **13-4** | TTT/**Azo**/TTT/**Azo/**TTT/**Azo/**TT/**Azo**/TTGTTTCTTTTTTTTTTTTATTTTAAAAAAAAAAAAA |
| **13-nt duplex control** | CTCCTCCTCCTCTCTTTGCTTTTTTTTTTTTTCATTTGAAAAAAAAAAAAA |
| **Apt-13-0** | /**$NH_2$**/CACCTGGGGGAGTATTGCGGAGGAAGGTTTAAAAAAAAAAAAATTTATTTTTTTTTTTTTCATTGTTTTTTTTTTTTTATATTCCCAGGTG-Cy3 |
| **Apt-13-3** | /$NH_2$/CACCTGGGGGAGTATTGCGGAGGAAGGTTTAAAAAAAAAAAAATTTATTTTTTTTTTTTTCATTGTTT/**Azo**/TTT/**Azo**/TTT/**Azo/**TTTTATATTCCCAGGTG-Cy3 |

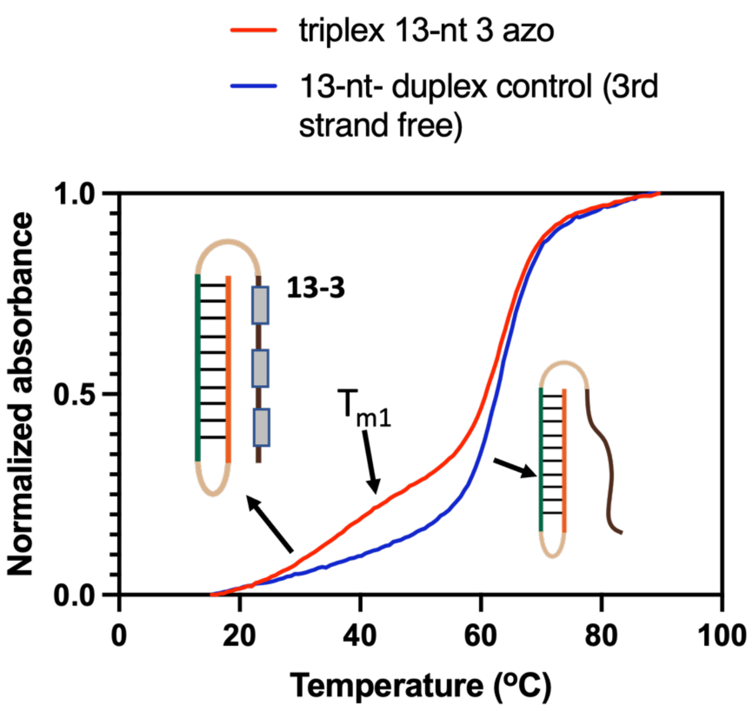


**Supporting Figure S1 | Melting curves for 13-3 and a control duplex.** $T_{m1}$ was not observed in the negative control, in which the third Hoogsteen strand has been mutated so that it does not form an intramolecular triplex.

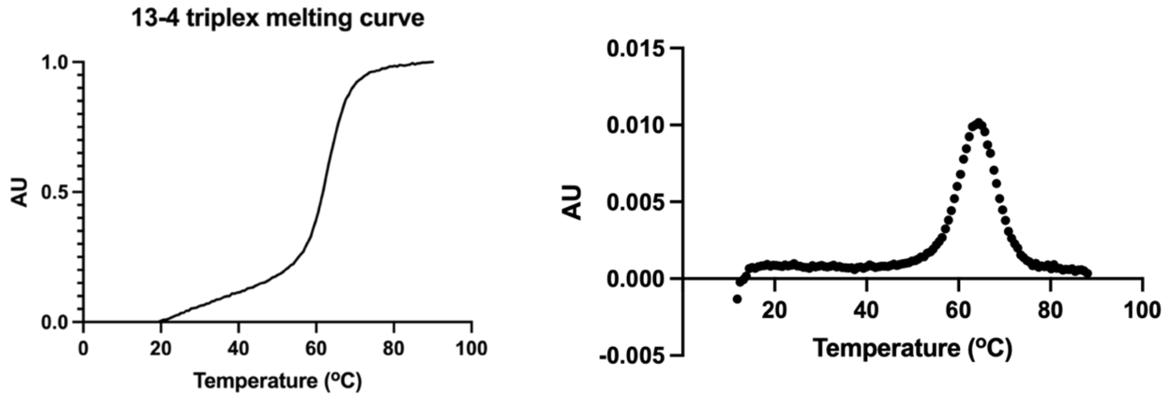


**Supporting Figure S2 | Melting curve of the 13-4 triplex**. Righthand panel shows first derivative of the melting curve. We observed no $T_{m1}$ for the **13-4** triplex structure.

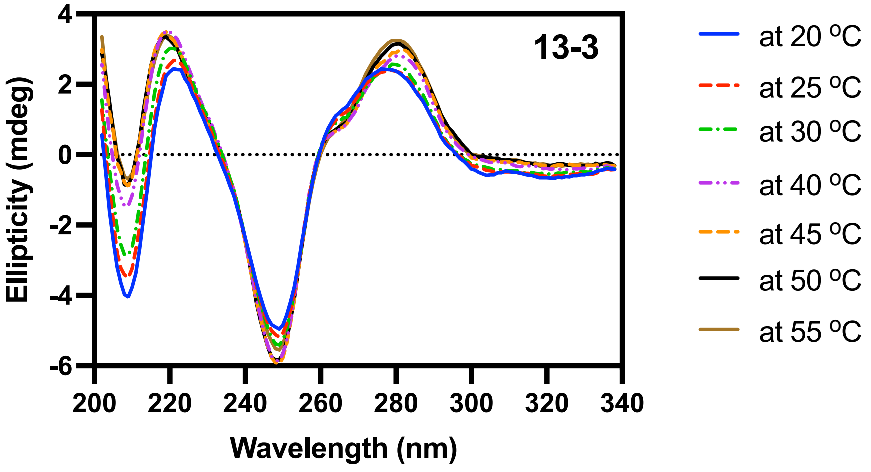


**Supporting Figure S3 | CD spectra of 13-3 at different temperatures**. As temperature increases, the Hoogsteen base-pairing gradually becomes weaker, and the peak around 210 nm (characteristic for triplexes) gradually decreases in amplitude.

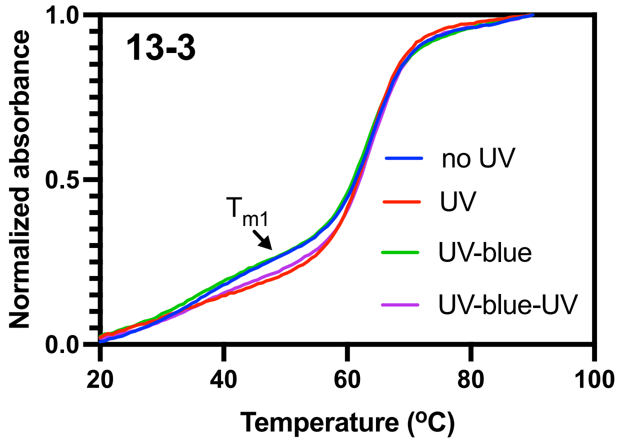


**Supporting Figure S4 | UV-Vis melting curves for 13-3 after several cycles of UV and blue light**.

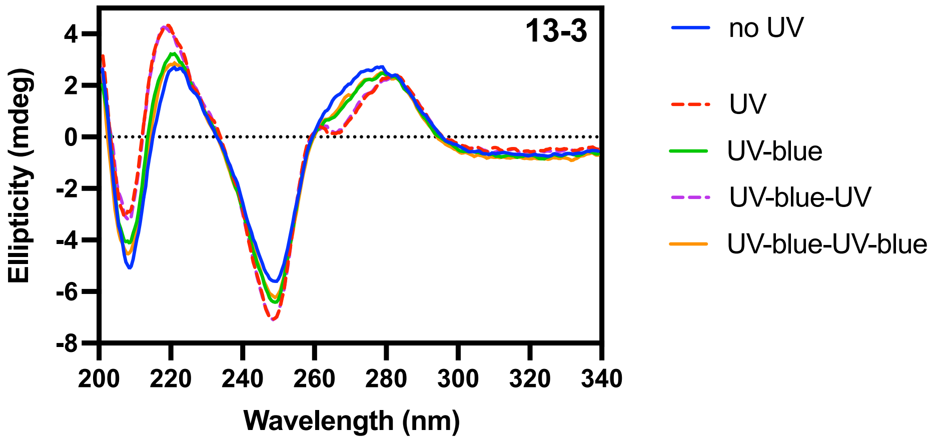


**Supporting Figure S5 | CD spectra for 13-3 after several cycles of UV (3 min) and blue light (1 min)**.

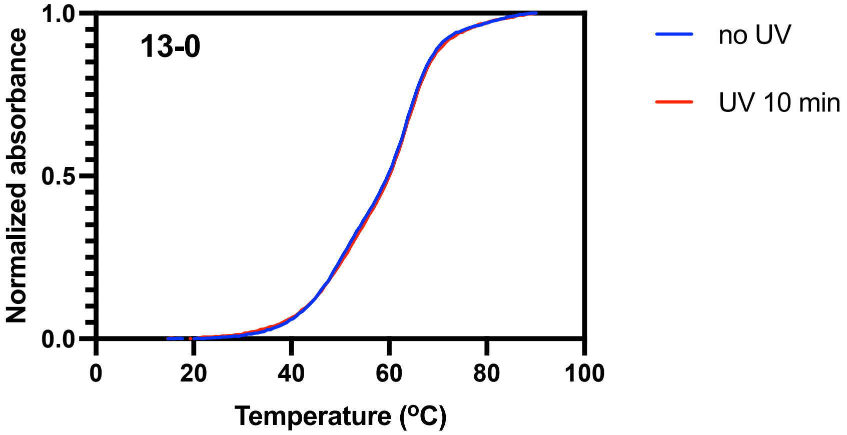


**Supporting Figure S6 | Melting curves of the 13-0 control before and after UV exposure in buffer**. The melting curve of **13-0** remains unchanged after 10 min UV exposure.

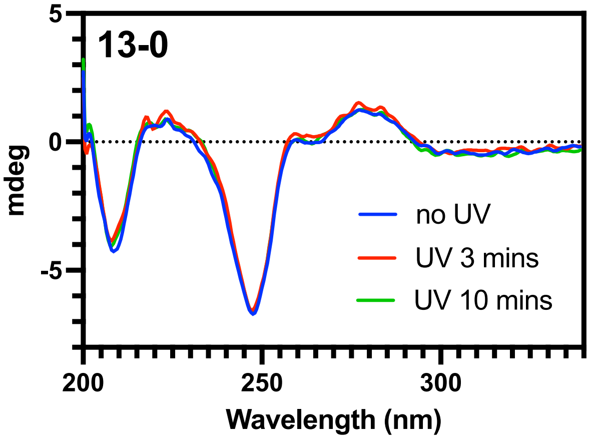


**Supporting Figure S7 | CD spectra of 13-0 after UV exposure**. **13-0** was exposed to 0-, 3-, or 10-min UV exposure. The CD traces showed that UV does not have any impact on azobenzene-free triplex behavior, as demonstrated by unchanged amplitude at 210 nm.

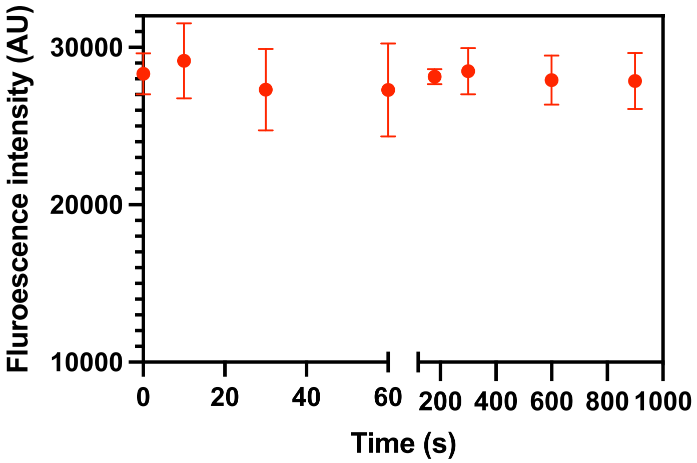


**Supporting Figure S8 | Photobleaching of a DNA-Atto 643 over the course of 15 mins**. A random DNA (sequence: 5' /$NH_2$/CATATATTGAGGAGTTATTA-3') was first functionalized with Atto 643-NHS. The Atto 643-DNA solution in 1x PBS/ 1mM $Mg^{2+}$ at 100 nM was then exposed to UV light (365 nm, intensity ~144-187 $mW/cm^2$) at different time points (0, 10 s, 30 s, 1 min, 3 mins, 5 mins, 10 mins and 15 mins). The samples were collected at each time point and were measured using a Synergy H1 microplate reading (BioTeK) in triplicate with Cy5 filter cube (Ex 625 nm/ Em 685). We observed no significant photobleaching occurred to the dye after 15 mins of UV exposure.

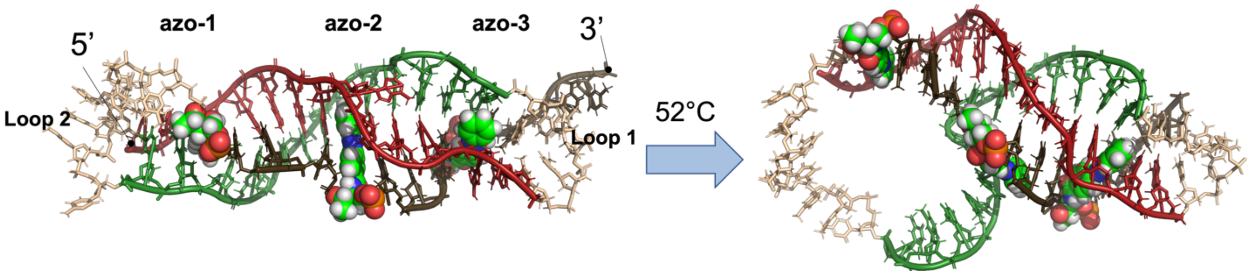


**Supporting Figure S9 | Comparison of the initial and final snapshots from the simulation of 13-3 triplex at 52 °C**. The DNA triplex and loops are shown in cartoon and stick representations, while the azobenzenes are represented as spheres. Unfolding near the loop 2 region and azo-1 in the triplex is shown at the end of the 500 ns simulation.

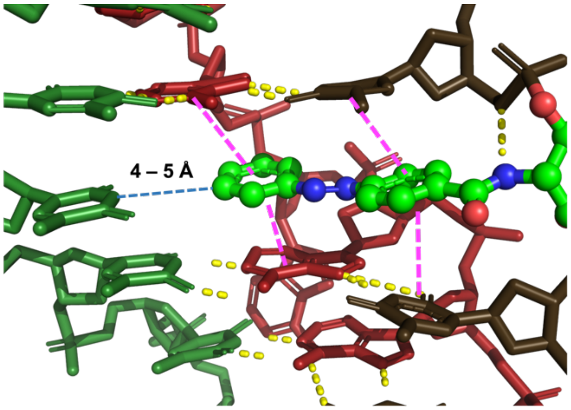


**Supporting Figure S10 | Illustration of the intercalation of azobenzene within neighboring bases in 13-3**. The DNA strands are shown as sticks and azo-2 is shown as balls and sticks, with hydrogen atoms hidden for clarity. Hydrogen bonds are shown as yellow dashes, and pi-pi stacking as magenta dashes. The closest distance between the azobenzene to the nearest nucleotide is 4–5 Å (side chain) and 7–25 Å (backbone).

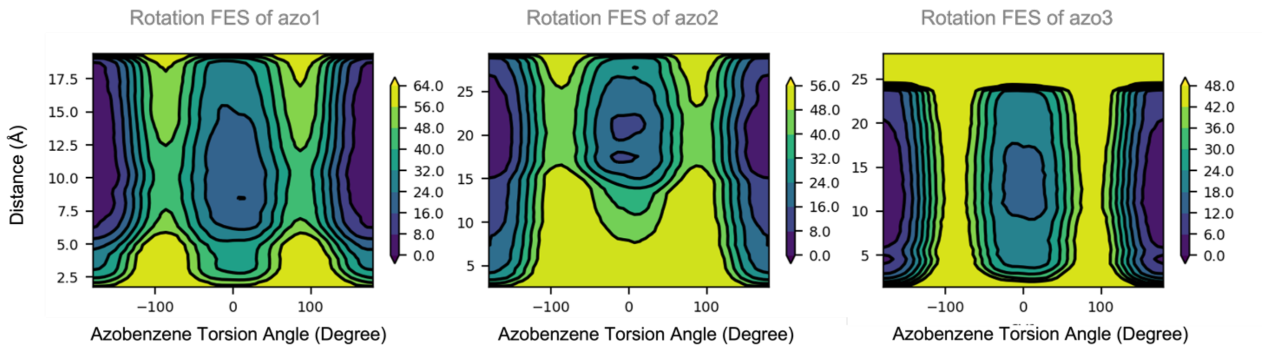


**Supporting Figure S11 | 2D metadynamics simulations of the 13-3 design for the rotation of each azobenzene in the folded triplex.** Charts show the free energy landscape under folding/unfolding and azobenzene rotation. The distance collective variable is defined as the distance from the azobenzene to the nearest first or second domain nucleotide backbone, while the angle distance is defined as the C-N=N-C dihedral in the azobenzene (*cis*- = 0°, *trans*- = 180° or -180°). Along the rotation of azobenzene from -180° to 0° (**Fig. 4b**), the second azobenzene (azo-2) had the highest energetic barrier (45.2 ± 2.1 kcal/mol) compared with the azo-1 (44.0 ± 1.4 kcal/mol) and azo-3 (42.9 ± 1.5 kcal/mol).
